# Nanobody mediated neutralization reveals an Achilles heel for norovirus

**DOI:** 10.1101/2020.02.07.938506

**Authors:** Anna D. Koromyslova, Jessica Michelle Devant, Turgay Kilic, Charles D. Sabin, Virginie Malak, Grant S. Hansman

## Abstract

Human norovirus frequently causes outbreaks of acute gastroenteritis. Although discovered more than five decades ago, antiviral development has, until recently, been hampered by the lack of a reliable human norovirus cell culture system. Nevertheless, a lot of pathogenesis studies were accomplished using murine norovirus (MNV), which can be grown routinely in cell culture. In this study, we analysed a sizeable library of Nanobodies that were raised against the murine norovirus virion with the main purpose of developing Nanobody-based inhibitors. We discovered two types of neutralizing Nanobodies and analysed the inhibition mechanisms using X-ray crystallography, cryo-EM, and cell culture techniques. The first type bound on the top region of the protruding (P) domain. Interestingly, the Nanobody binding region closely overlapped the MNV receptor-binding site and collectively shared numerous P domain-binding residues. In addition, we showed that these Nanobodies competed with the soluble receptor and this action blocked virion attachment to cultured cells. The second type bound at a dimeric interface on the lower side of the P dimer. We discovered that these Nanobodies disrupted a structural change in the capsid associated with binding co-factors (i.e., metal cations/bile acid). Indeed, we found that capsids underwent major conformational changes following addition of Mg^2+^ or Ca^2+^. Ultimately, these Nanobodies directly obstructed a structural modification reserved for a post-receptor attachment stage. Altogether, our new data show that Nanobody-based inhibition could occur by blocking functional and structural capsid properties.

**AUTHOR SUMMARY:** This research discovered and analysed two different types of MNV neutralizing Nanobodies. The top-binding Nanobodies sterically inhibited the receptor-binding site, whereas the dimeric-binding Nanobodies interfered with a structural modification associated with co-factor binding. Moreover, we found that the capsid contained a number of vulnerable regions that were essential for viral replication. In fact, the capsid appeared to be organized in a state of flux, which could be important for co-factor/receptor binding functions. Blocking these capsid-binding events with Nanobodies directly inhibited essential capsid functions. Moreover, a number of MNV-specific Nanobody binding epitopes were comparable to human norovirus-specific Nanobody inhibitors. Therefore, this additional structural and inhibition information could be further exploited in the development of human norovirus antivirals.

## INTRODUCTION

Norovirus belongs to the *Caliciviridae* family of non-enveloped, single-stranded positive-sensed RNA viruses [1]. The *Norovirus* genus comprises at least seven genogroups (GI-GVII), where GI, GII, and GIV infect humans [2]. Worldwide, human norovirus is one of the leading causes of outbreaks of acute gastroenteritis [3–5]. There are still no antivirals or vaccines for norovirus. Moreover, clinical trials with norovirus virus-like particle (VLP) vaccines have had limited success [6–8].

Caliciviruses also infect other animals and include rabbit haemorrhagic disease virus (RHDV), feline calicivirus (FCV), and murine norovirus (MNV). Pathogenic studies using MNV have provided an abundance of neutralization, vaccine development, and pathogenesis information, since MNV is grown routinely in cell culture and a reliable reverse genetics system is available [9–11].

Structural studies have shown that the virion capsid (VP1) has a T=3 icosahedral symmetry. The capsid comprises of 180 VP1 copies, which are structured in three quasi-equivalent subunits that fold into A/B and C/C dimers [12]. VP1 can be divided into two domains, termed shell (S) and protruding (P) domain. The S domain forms the inner core and surrounds the viral RNA. The P domain forms protruding spikes and contains the main determinants for host binding factors, which can include binding sites for histo-blood group antigens (HBGAs), bile acids, bivalent metal cations, and the receptor CD300lf [13–17]. The P domain is further divided into two subdomains, a distal P2 subdomain, and a lower P1 subdomain that is connected to the S domain via a flexible hinge region [12, 18–20].

Unfortunately, limited research is focused on the discovery of norovirus antivirals. The different steps in the replication cycle, including cell attachment and entry, replication and translation, and virion assembly offer many ideal targets. A lot of antiviral development is targeted against the capsid, especially regions that bind co-factors. Recent studies discovered human norovirus-specific monoclonal antibodies (mAbs) and Nanobodies that sterically blocked the HBGA pocket [21–26]. Other studies using MNV also showed that blocking the MNV CD300lf receptor-binding pocket with MNV-specific mAbs inhibited viral replication [27, 28].

In this study, we screened a large library of MNV-specific Nanobodies in order to identify Nanobody-based inhibitors. We found several candidates that block replication and the structural basis of neutralization was analysed. Overall, our findings exposed crucial roles of capsid conformational modifications and described two Nanobody-based inhibition mechanisms.

## RESULTS

### Nanobody neutralizing and binding capacities

A library of 58 MNV virion-specific Nanobodies was analysed in order to identify candidates that presented superior neutralizing and binding properties. A total of 51 distinct Nanobody families (based on CDR sequence diversity) were produced and analysed. In an attachment assay, most Nanobodies reduced the number of MNV plaques, where 38 had greater than 70% inhibition at 20 μg/ml (Fig. S1A). Fifteen Nanobodies reduced the number of plaques by more than 75% at 2 μg/ml (Fig. 1). Dilution inhibition curves of these Nanobodies yielded IC_50_ values ranging between 0.03 to 1.6 μg/ml, where NB-5867 and NB-5894 were the most effective (IC_50_ = 0.03 and 0.09 μg/ml, respectively) (Fig. S1B). ELISA data showed that these Nanobodies bound strongly to virions, having cut-off concentrations ranging between 0.9 and 4.4 ng/ml.

**Figure 1.**
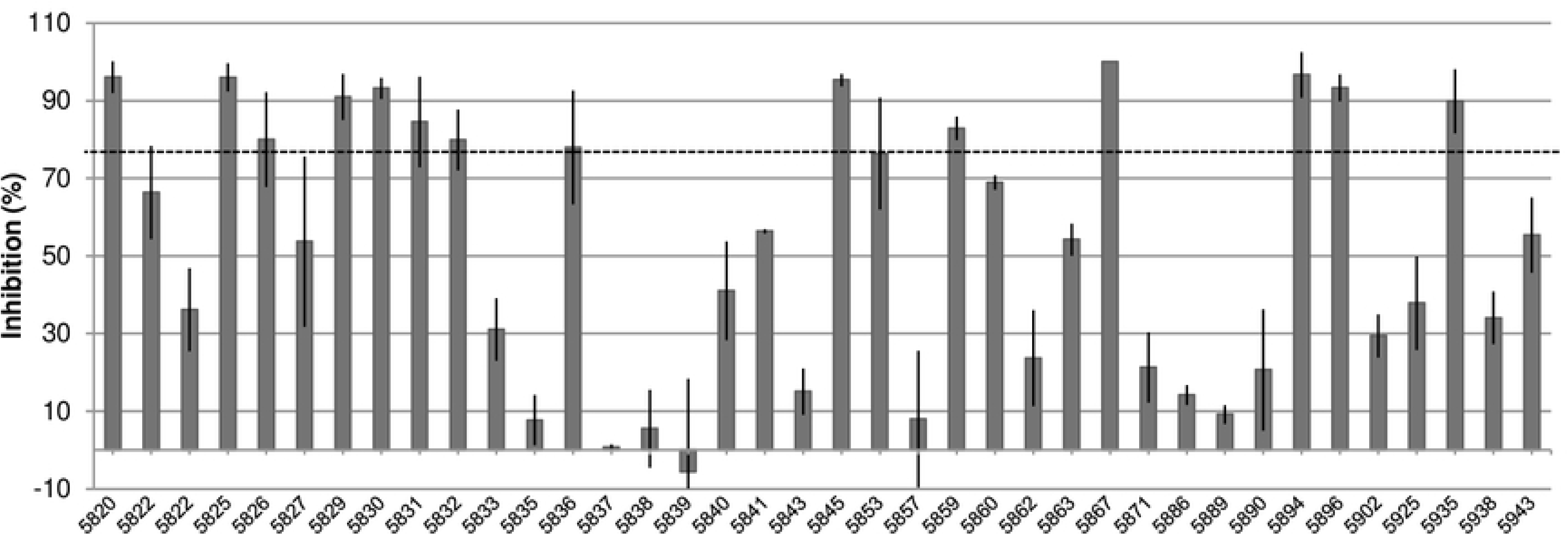
Neutralization assay. The neutralization capacity of MNV-specific Nanobodies was evaluated using a plaque assay (attachment assay). At 2 μg/ml, fifteen Nanobodies showed more than 70% inhibition (dashed line). All experiments were performed in triplicates and standard deviation is shown.

The binding properties for eight Nanobodies were further analysed using ITC and K_d_ values ranged between 0.03 to 13.9 nM (Table S1). The binding reactions were exothermic and were fitted into a one-site binding model (stoichiometry ∼1). However, entropy contribution was variable, even between Nanobodies from the same family, and ranged between −33 and 32 KJ/mol. Overall, these neutralizing and binding results revealed that several Nanobodies had high affinities and excellent neutralization capacities.

### X-ray crystal structures of P domain and Nanobody complexes

In order to show how neutralizing Nanobodies bound to the capsid, the X-ray crystal structures of MNV P domain and Nanobody complexes were determined (Table S2). The electron densities of the P domain and Nanobody complexes were well resolved and water molecules were observed in all structures. The overall structure of the P domain in all complex structures was highly similar to the apo P domain, except for several loop movements. All Nanobodies had the typical immunoglobulin fold and the CDRs primarily interacted with the P domain. Two Nanobodies (NB-5853 and NB-5867) bound on the top of the P2 subdomain and two Nanobodies (NB-5820 and NB-5829) bound at the dimeric interface on the side of the P1 subdomain (Fig. 2).

**Figure 2.**
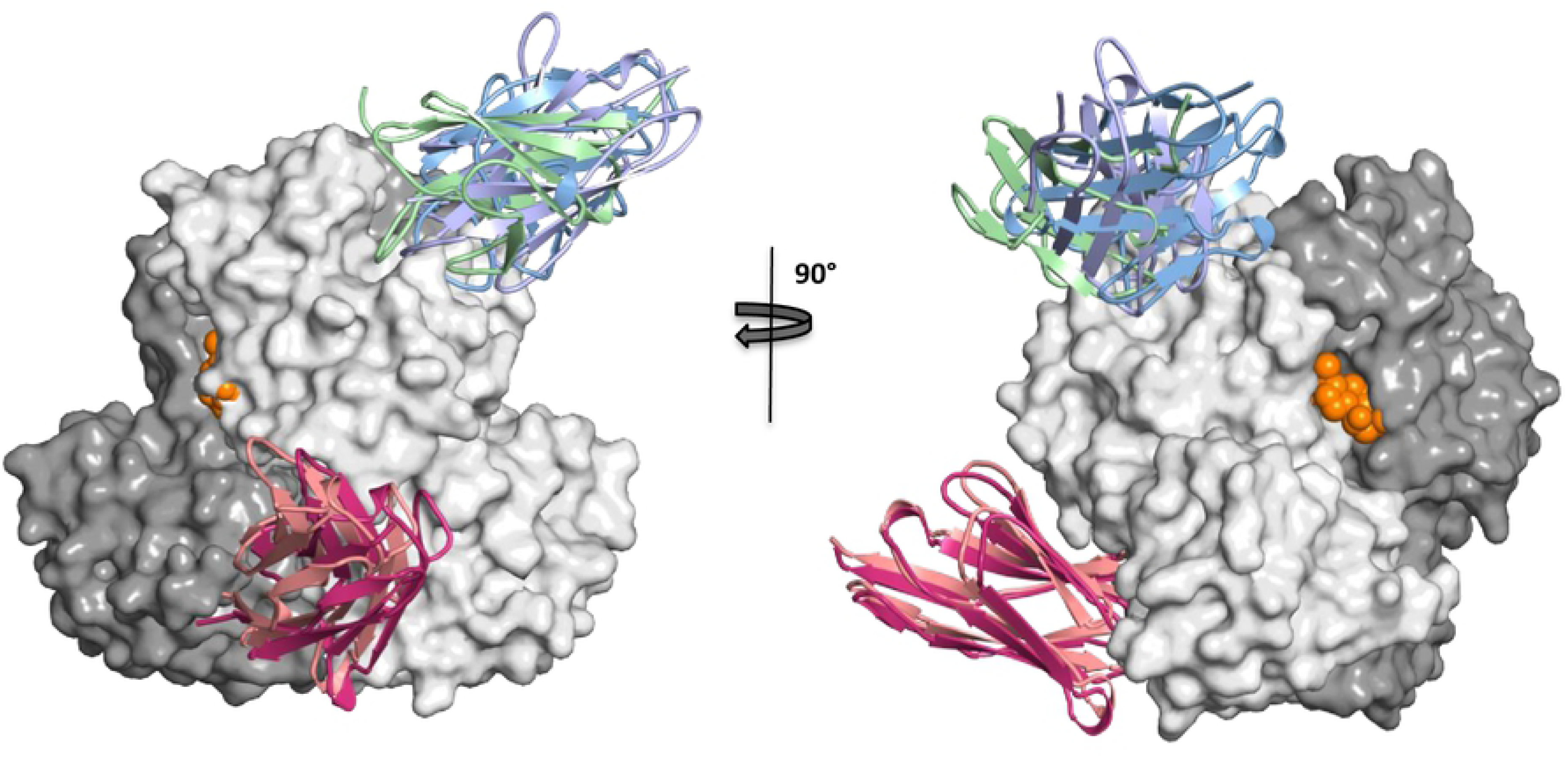
Nanobody binding sites on the P dimer. The X-ray crystal structures of the P domain Nanobody complexes were superimposed onto one P dimer in order to show all four Nanobody binding sites. Nanobodies were colored accordingly, NB-5853 (sky), NB-5867 (purple), NB-5820 (salmon), and NB-5829 (warm pink). NB-5853 and NB-5867 bound on the top of the P2 subdomain, whereas NB-5820 and NB-5829 bound on the side of the P1 subdomain and at a dimeric interface. The MNV CD300lf (green) receptor and bile acid (orange) were superpositioned onto the P dimer for comparison.

### Structure of MNV P domain and NB-5853 complex

The X-ray crystal structure of P domain NB-5853 complex was solved to 1.96 Å resolution. A network of hydrogen bonds was formed between the P domain and NB-5853 (Fig. 3A and Table S3). The majority of NB-5853 binding residues were located in CDR3. Five P domain residues (Q334, G400, E356, T362, and N364) formed eight direct hydrogen bonds with NB-5853 residues. Furthermore, five P domain residues (T301, V304, A365, F375, and Y399) were involved in hydrophobic interactions with NB-5853 residues. Additionally, a number of water-mediated bonds were formed with the P domain and NB-585. Overall, these findings showed that the Nanobody was held tightly by one P domain monomer.

**Figure 3.**
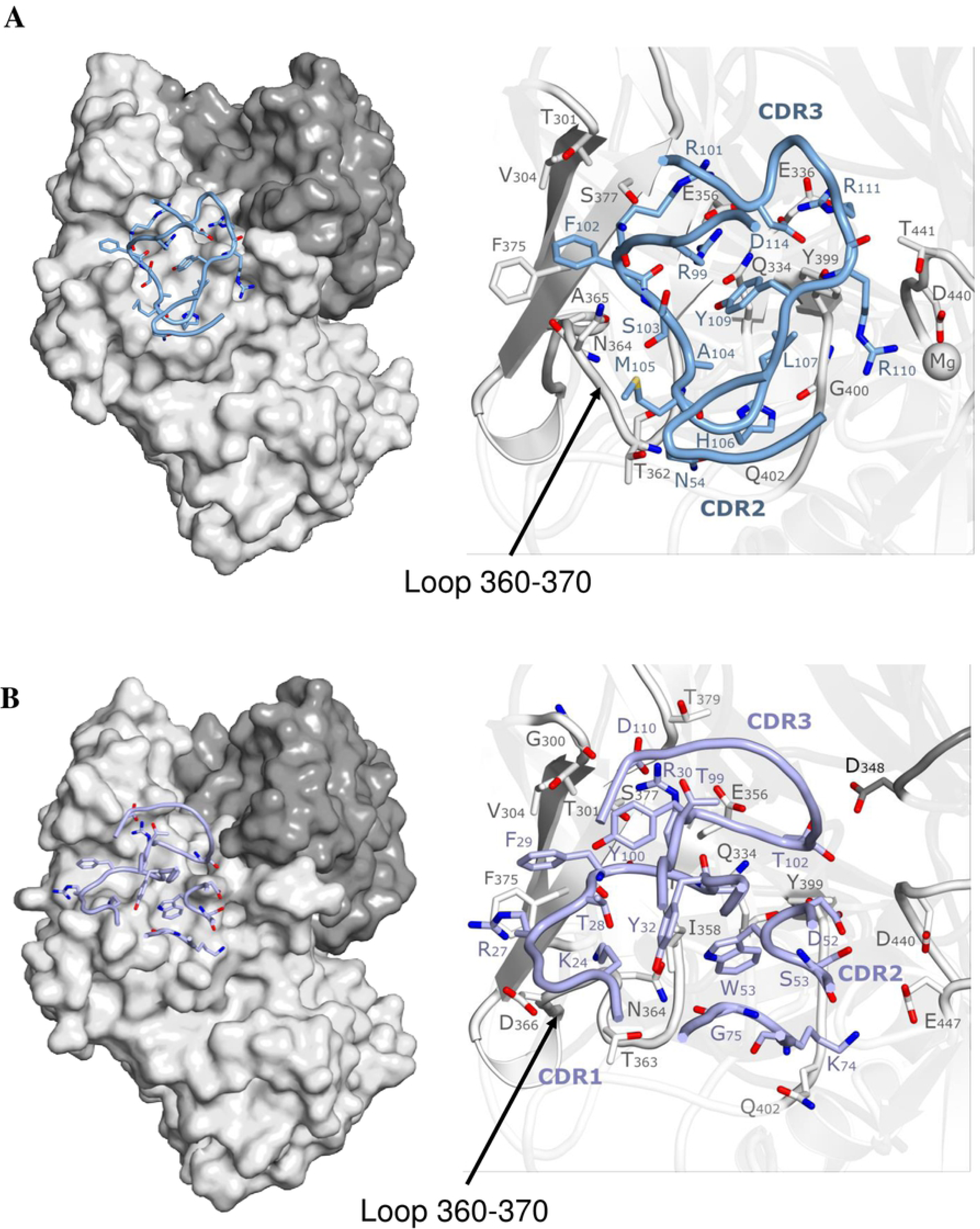
Interaction of the top-binding Nanobodies. These Nanobodies interacted with one P domain and closely overlapped the CD300lf binding site (see Fig. S2). Hydrogen bond distances were cut at 3.3 Å, though the majority were ∼2.8 Å. (A) NB-5853 was held with network of P domain direct hydrogen bonds and hydrophobic interactions (Nanobody residues binding residues shown for simplicity). NB-5853 directly interacted with several P domain residues on the loop 360-370 (i.e., T362, N364, and A365). (B) NB-5867 interacted with both P domains, although most bonds were provided by one monomer. NB-5867 also interacted with several P domain residues on the loop 360-370 (T363 and N364).

Remarkably, the NB-5853 binding site closely overlapped the MNV CD300lf receptor footprint (Fig. S2). In fact, NB-5853 interacted with nine P domain residues that held CD300lf (T301, V304, Q334, E356, N364, F375, S377, Y399, and I358). Moreover, NB-5853 binding resulted in a P domain conformational change comparable to binding soluble CD300lf i.e., in loops covering residues 341-351 and 360-370. One of these loops (341-351) was “closed” in apo P domains and “open” when CD300lf was bound [13, 29]. These findings indicated that NB-5853 bound to the P domain similarly as the soluble receptor. Moreover, these results suggest that NB-5853 might directly interfere with CD300lf binding.

### Structure of MNV P domain and NB-5867 complex

The X-ray crystal structure of the P domain in complex with NB-5867 was solved to 2.19 Å resolution. The binding site of NB-5867 was closely similar to NB-5853 (Fig. S2), except for an additional hydrogen bond provided by a residue (D348) on the other monomer (Table S3). All three CDRs of NB-5867 were involved in binding (Fig. 3B) Eight P domain residues (T301, Q334, T363, N364, S377, T379, Y399, and D348^monomer2^) formed fourteen direct hydrogen bonds with NB-5867 residues. Five P domain residues (T301, V304, I358, F375, and Y399) were involved in hydrophobic interactions with NB-5867 residues and a number of water-mediated bonds provided additional interactions between the two proteins. In summary, our results showed that NB-5867 was tightly held by mostly one P monomer.

NB-5867 interacted with nine P domain residues that also bound CD300lf (T301, V304, Q334, E356, I358, N364, F375, S377, and Y399) (Fig. S2). Moreover, the loops covering residues 341-351 and 360-370 shifted into an equivalent position as in the NB-5853 and CD300lf complexes. Overall, these findings showed that the top-binding Nanobodies overlapped the CD300lf binding site and inhibition could interfere with receptor binding events.

### NB-5853 and NB-5867 inhibition mechanism

In order to better understand the inhibition mechanism of the top-binding Nanobodies, a series of competitive ITC measurements using MNV P domain, sCD300lf, bile acid (GCDCA), and CaCl_2_ was performed (Fig. 4A). The binding of NB-5867 to the P domain was not affected by the addition of GCDCA or CaCl_2_, although the binding affinity in presence of GCDCA was lower than with CaCl_2_ and PBS (K_d_ = 143 nM vs K_d_ = 14 and 23 nM, respectively). As expected, when the P domain was pre-incubated with NB-5867, sCD300lf did not bind to the P domain (Fig. 5A). Consequently, the ITC data confirmed the structural findings that these Nanobodies blocked the receptor-binding pocket.

**Figure 4.**
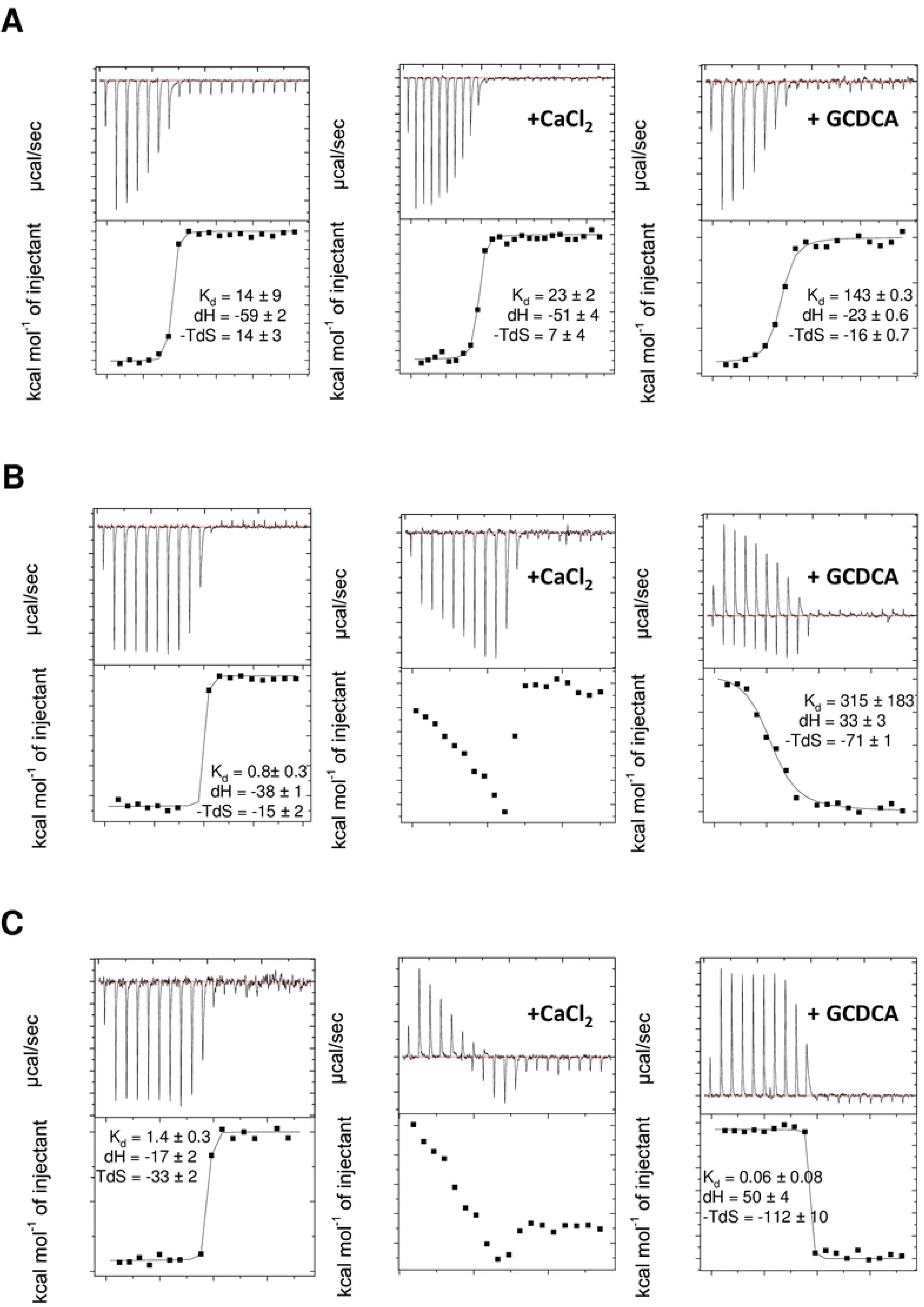
Bile acid and CaCl_2_ influence on the thermodynamic properties of Nanobody binding. Nanobodies were titrated into P domain in presence of PBS (control), 5 mM CaCl_2_, and 50 µM GCDCA. Affinity values (K_d_) are presented in nM, whereas enthalpy (dH) and entropy (-TdS) are measured in KJ/mol. All experiments were performed twice. (A) NB-5867 binding affinity was not affected by CaCl_2_, but reduced with GCDCA by 6-10 times (from 14 nM to 143 nM). (B) NB-5820 enthalpy and entropy were both influenced by CaCl_2_ and GCDCA. For GCDCA, the enthalpy input was reversed from −38 KJ/mol to 33 KJ/mol and entropy input changed from −15 KJ/mol to −71 KJ/mol. (C) NB-5829 properties were similar to NB-5820, where the reversed enthalpy increased (−17 KJ/mol to 50 KJ/mol) and a change in entropy was measured (−33 KJ/mol to −112 KJ/mol) when GCDCA was added. For CaCl_2_ no binding model could be fitted to the data for either NB-5820 or NB-5829.

**Figure 5.**
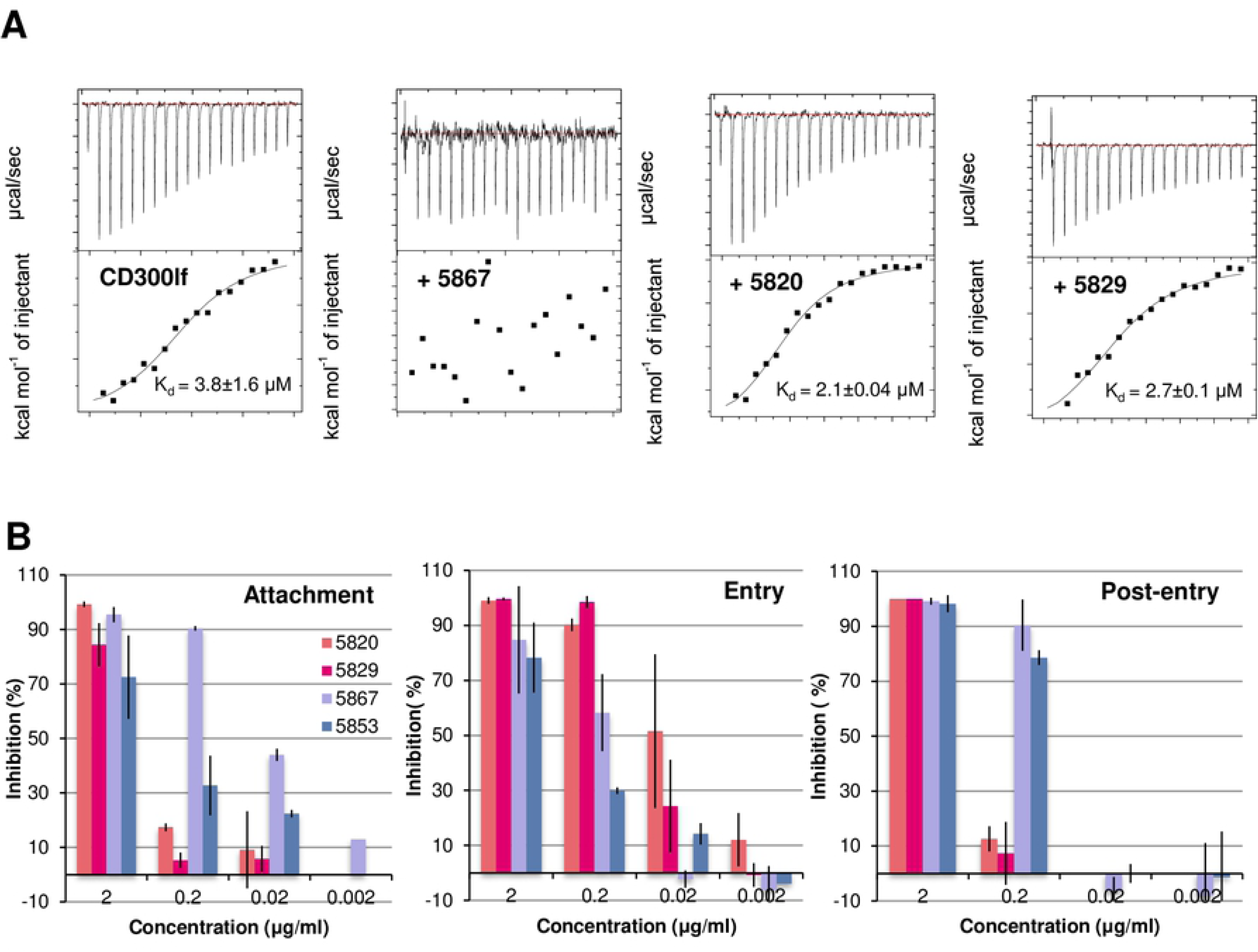
Nanobody competition with sCD300lf and functional assays. The ability of Nanobodies to complete with CD300lf were analysed using ITC. (A) sCD300lf was titrated into P domain supplemented with CaCl_2_ and GCDCA. The binding reaction was exothermic with K_d_ = 3.8 ± 1.6 μM. Next, the P domain was pre-mixed with CaCl_2_, GCDCA, and Nanobodies NB-5820, NB-5829, or NB-5867 in a 1:2 molar ratio, followed by sCD300lf titration. NB-5820 and NB-5829 did not change sCD300lf-binding characteristics and the affinity with or without Nanobody was similar. For NB-5867, no binding signal was observed, which indicated that this Nanobody competed with sCD300lf. (B) Nanobody inhibition of MNV infection was analysed using at the attachment, entry, and post-entry stages. Nanobodies binding on the top were most effective in attachment and post-entry stage, whereas Nanobodies binding at the dimeric interface blocked the entry (post-attachment) stage of MNV infection. All experiments were performed in triplicates and standard deviation was shown.

In order to further identify other stage(s) of the inhibition, a series of modified entry (post-attachment) and post-entry assays using the MNV cell culture system were performed (Fig. 5B). For the entry (post-attachment) assay, the virions were first allowed to attach to the cell surface for 3 h at 4°C, then unbound virus was removed and Nanobodies were added to the culture medium for 1 h (viral infection stage). After infection, cells were washed, overlaid with agarose, and the number of plaques was analysed at 2 dpi. For the post-entry assay, the Nanobodies were only added after MNV infection and were present in agarose overlay until the number of plaques was analysed. NB-5867 showed strong inhibition in attachment (IC_50_ = 0.02 μg/ml) and post-entry (IC_50_ = 0.06 μg/ml). The inhibition ability was lower in the entry assay (IC_50_ = 0.16). Altogether, these cell culture results showed that NB-5867 effectively blocked MNV at the cell attachment stage.

### Structure of MNV P domain and NB-5820 complex

The X-ray crystal structure of the MNV P domain NB-5820 complex was determined to 1.72 Å resolution. The P domain interactions were formed between all CDRs and the C-terminus of NB-5820 (Fig. 6A and Table S3). Four P domain residues (V234, Q236, L481; and D272^monomer2^) formed four direct hydrogen bonds with NB-5820 residues. One P domain residue (Val234) was involved in one hydrophobic interaction with NB-5820. Thirteen water-mediated bonds were also observed. For NB-5820, the P domain loop covering residues 360-370 was observed in a comparable position as with NB-5853, NB-5867, and CD300lf. Conversely, the loop covering residues 341-351 was in a closed position (unlike NB-5853, NB-5867, and CD300lf).

**Figure 6.**
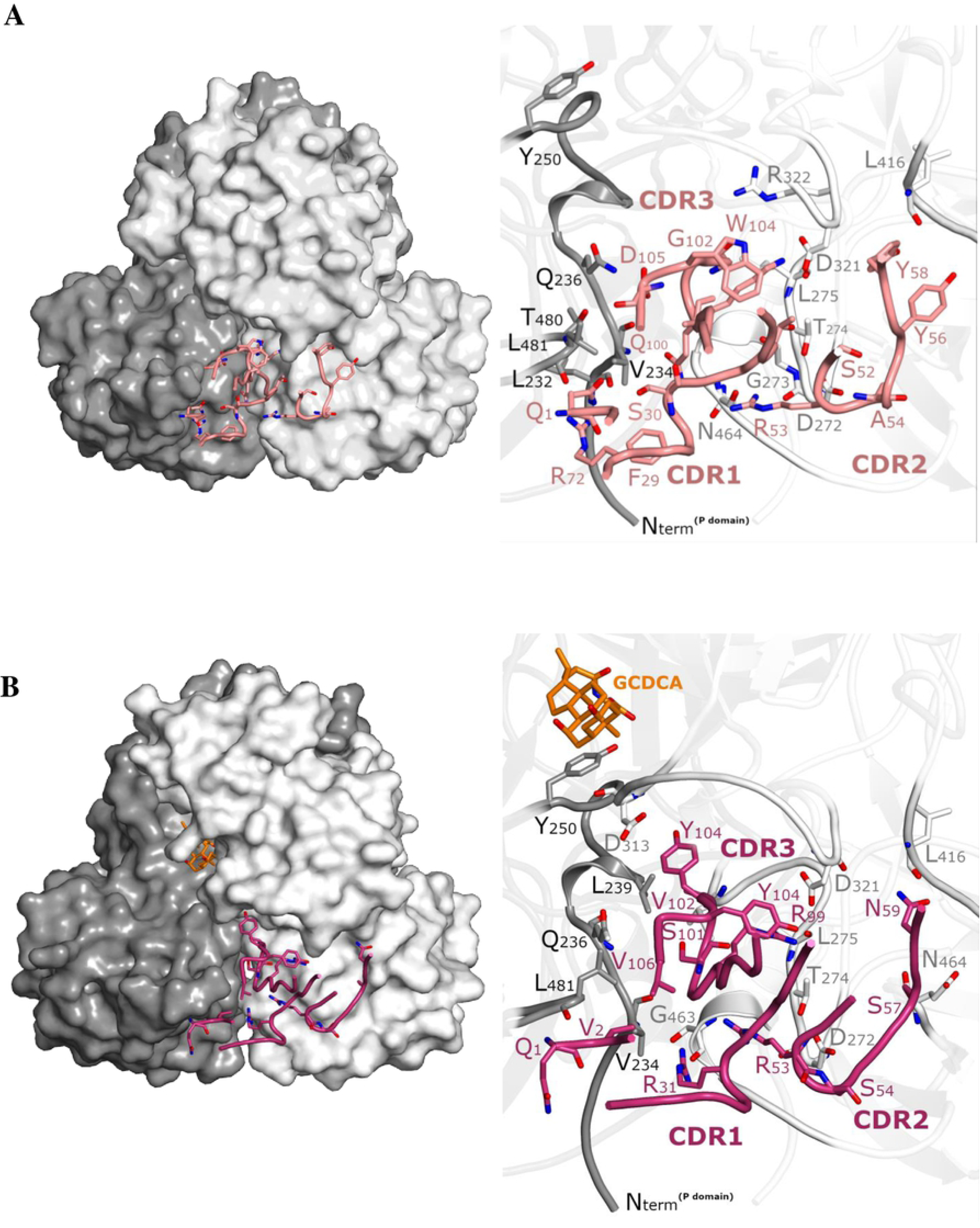
Interaction of the dimeric-binding Nanobodies. Both Nanobodies bound on the P dimer with a similar footprint. Hydrogen bond distances were cut to 3.3 Å. (A) NB-5820 was mostly held with residues from one P domain monomer. (B) Residues from both P domain monomers firmly held NB-5829 with direct hydrogen bonds and hydrophobic interactions.

Interestingly, NB5820 CDR1 and CDR3 interacted with several residues (Val234 and Gln236) located between residues 228-250 of the P1 subdomain (Fig. 6A). This stretch of residues contained the hinge region that connected the S and P1 subdomains, while Tyr250 aided the binding of bile acid. This initial observation suggested that NB-5820 might influence the hinge region and/or bile acid-binding function(s).

### Structure of MNV P domain and NB-5829 complex with bile acid

Unlike NB-5820, our attempts to produce crystals for MNV P domain and NB-5829 complex were unsuccessful. However, when bile acid (GCDCA) was added to the complex solution, we obtained crystals that diffracted to 2.15 Å resolution. The complex structure was found to also contain GCDCA and Mg^2+^. Bile acid formed a number of hydrogen bonds with the P domain (R390, R392, R437, and W245) as well as hydrophobic interactions (A247, Y250, A290, G314, Q340, Y435, and M436). These binding interactions were almost identical to a previously released MNV P domain GCDCA complex structure [14].

NB-5829 bound at the similar P domain dimeric interface as NB-5820 (Fig. S2). However, all three NB-5829 CDRs interacted with the P domain. Also, the total number of interactions was greater than for NB-5820 (Fig. 6B and Table S3). Eight P domain residues (D272, T274, L275, D313, D321, A462, S463, and E494) formed ten direct hydrogen bonds with NB-5829 residues. Three P domain residues (I281, L239^monomer2^, and L481^monomer2^) were involved in five hydrophobic interactions and water-mediated bonds provided additional connections between the two proteins.

Another interesting feature of NB-5829 binding was the positions of loops covering residues 341-351 and 360-370. For NB-5829, these loops were positioned as observed for NB-5853, NB-5867, and CD300lf, but dissimilar to NB-5820 that positioned loop 341-351 in the closed position (Fig. 6B). This finding suggested that loop 341-351 could be opened for bile acid binding, but was closed for binding of dimeric-binding Nanobodies alone. More importantly, NB-5829 interacted with residue L239, which was located close to the hinge region. This finding suggested that NB-5829 and NB-5820 might have similar inhibition mechanisms.

### NB-5820 and NB-5829 inhibition mechanism

In previous studies with human norovirus, we showed that a broadly reactive Nanobody (Nano-26) bound on the side of the P domain and inhibited VLPs from binding to HBGAs [21, 30, 31]. We discovered that VLP aggregation and disassembly followed when mixed with Nano-26. Surprisingly, the Nano-26 binding site was remarkably similar to NB-5820 and NB-5829. Thus, we first suspected that NB-5820 and NB-5829 might have similar effects on MNV virions. However, when virions were treated with either these Nanobodies, neither disassembly nor particle aggregation occurred (Fig. S3). This finding suggested that NB-5820 and NB-5829 neutralization mechanisms were distinct from Nano-26.

Based on NB-5820 and NB-5829 binding sites, we also assumed that these Nanobodies might indirectly interfere with the receptor or co-factor binding functions. Therefore, a series of competitive ITC measurements using MNV P domain, sCD300lf, GCDCA, and CaCl_2_ were performed. The affinity of MNV P domain binding to sCD300lf (K_d_ = 4 μM) was not affected by the addition of NB-5820 and NB-5829 (K_d_ = 2 μM and 3 μM, respectively), indicating that there was no interference with the receptor binding (Fig. 5A). When NB-5829 was titrated into the P domain in the presence of GCDCA, the binding profile changed from exothermic to endothermic, reversing the enthalpy contribution (Fig. 4) and increasing entropy input with a resulting lower K_d_ = 0.06 nM.

For the NB-5820, the K_d_ value in the presence of GCDCA was lower than the P domain alone (315 nM vs. 0.8 nM). In the presence of CaCl_2_ the titration curves of NB-5829 and NB-5820 binding to the P domain showed a combination of exothermic and endothermic binding events that could not be fitted to a standard binding model. Based on these findings, our data suggested that bile acid and CaCl_2_ binding triggered long-distance conformational changes that in turn influenced Nanobody binding.

Following this result, we performed entry- (post-attachment) and post-entry inhibition assays with NB-5820 and NB-5829 (Fig. 5B). We found that NB-5820 and NB-5829 had similar blocking activities in attachment and post-entry assays (IC_50_ value = 0.5 - 0.9 μg/ml). Surprisingly, the inhibition in the entry assay was > 50 times higher (IC_50_ = 0.02 - 0.04 μg/ml). This data indicated that these Nanobodies mainly interfered with a post receptor binding stage.

### Cryo-EM structure of Nanobody binding to MNV virions

We first determined the cryo-EM structure of apo MNV virion at 4.6 Å resolution (Figs. S4 and S5A). The virion closely resembled previously solved structures, where the P dimers were raised off the shell by ∼15 Å [27]. After fitting in the P domain/NB-5829 complex structure into this virion structure we found that neighbouring Nanobodies clashed (Fig. S5B). This immediate result suggested that NB-5829 did not initially bind all 180 epitopes or a structural modification was necessary for complete Nanobody occupancy.

Next, the cryo-EM structure of MNV virion NB-5829 complex was solved to 4.7 Å resolution (Fig. S4). Similar to the apo virion, the complex virion had T=3 icosahedral symmetry and the P dimers were raised off the shell (Fig. 7A). This structure showed two additional densities per P dimer that corresponded to bound Nanobodies (Fig. 7B). In this structure, the Nanobodies did not clash with neighbouring P dimers or Nanobodies. Interestingly, NB-5829 caused the P dimers to rotate ∼42° clockwise. In this orientation, the Nanobodies pointed towards the centre of the 3- and 5-fold axes. This rotation likely allowed the Nanobodies to bind all possible epitopes.

**Figure 7.**
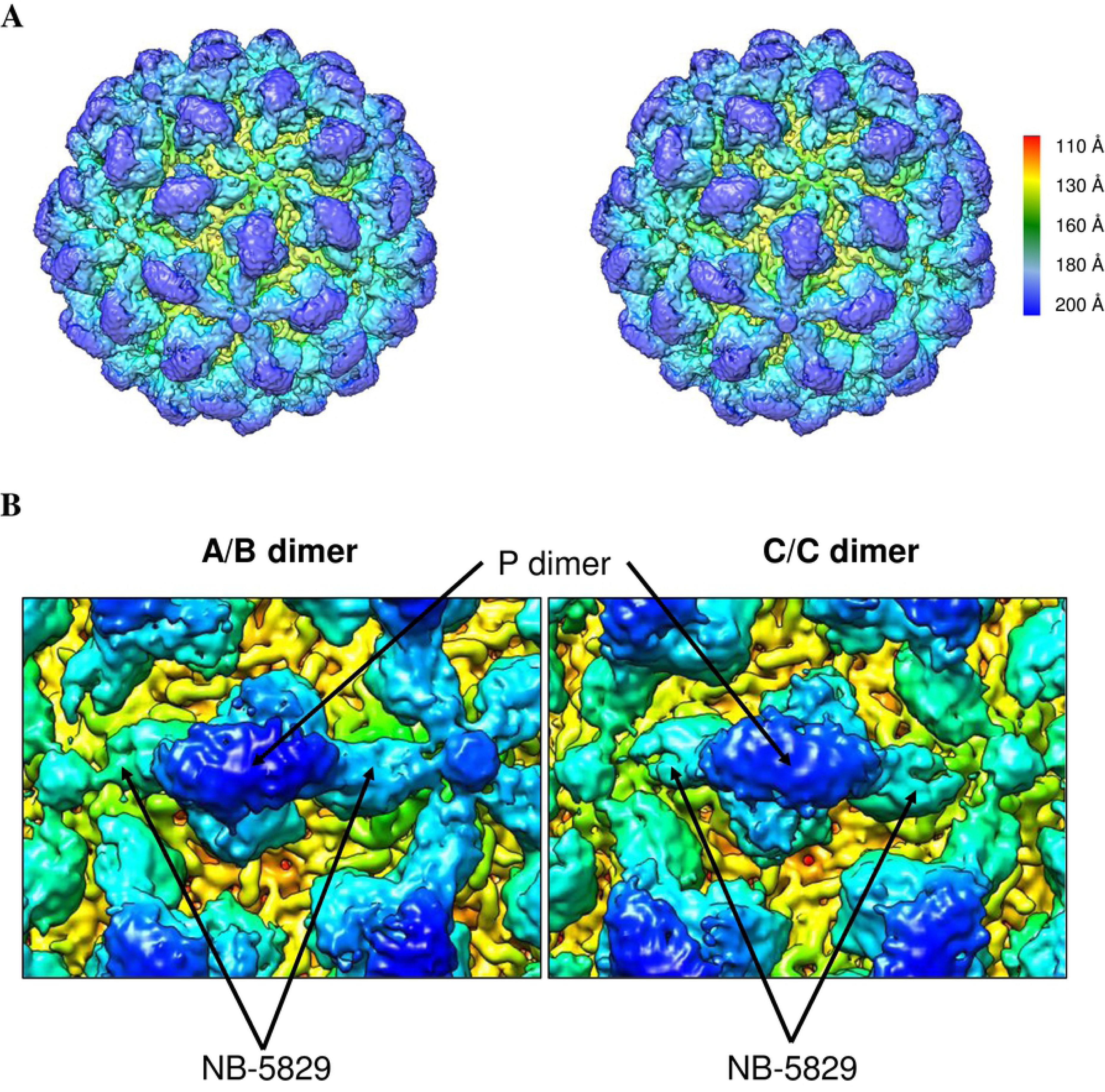
Cryo-EM structure of the MNV virion and NB-5829 complex. Icosahedral reconstruction of the virion and NB-5829 complex was solved to 4.7 Å resolution. (A) A stereo view of the complex structure colored by radius. (B) A close-up view of the complex, where NB-5829 was represented by additional density on the P domain. In this orientation, where NB-5829 pointed towards the centre of the 3- and 5-fold axes, all possible epitopes were occupied.

Structural refinement indicated that the S domain was better resolved than the P domain (8 Å resolution vs. 4 Å resolution, respectively). Therefore, focused reconstruction on the P dimers was performed to resolve the heterogeneity of the P domains. The P domain and NB-5829 complex structure was fitted into these P dimers and revealed that the P dimers occupied a variety of tilted positions (Fig. 8A). The A/B dimers were tilted up to ∼31° and the C/C dimers were tilted up to ∼34° (Fig. 8B). These results showed that the P dimers were indeed flexible (in terms of rotation and tilt). Moreover, the dimeric-binding Nanobodies forced these major structural changes, which may or may not be structurally detrimental for the virion.

**Figure 8.**
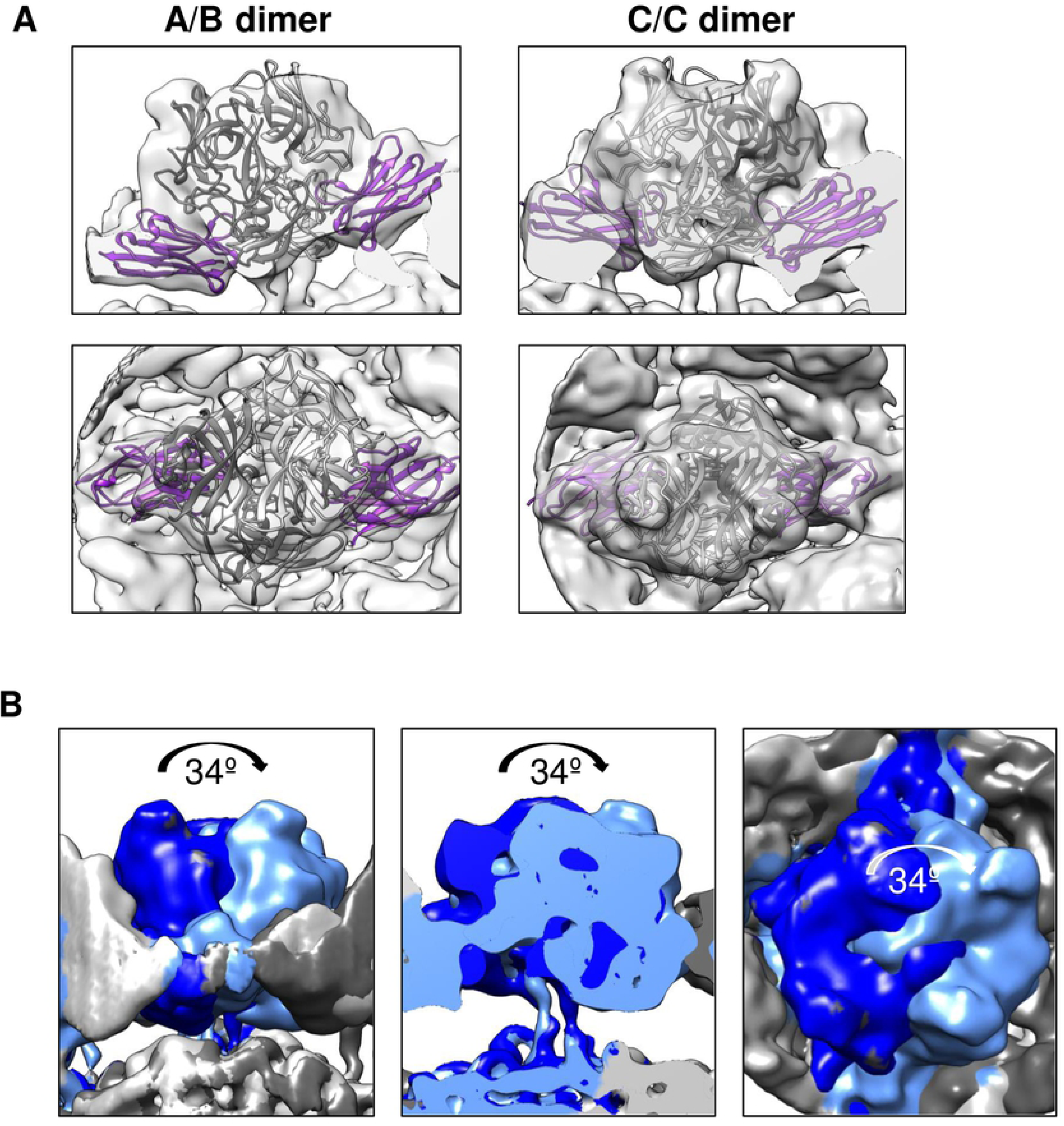
Focus reconstruction of P dimers. (A) Representative class of A/B and C/C dimers after focused reconstruction, fitted with the X-ray crystal structure of the complex (NB-5829: purple; and P domain: grey). Side view (upper panels) and top view (lower panels) showed that NB-5829 from the X-ray crystal structure fitted well into the model. (B) Focused reconstruction refinement also showed that the P dimers tilted up to ∼34° in C/C dimers. A surface representation of the P domain monomer rotation is shown moving from cyan to blue.

### Cryo-EM structure of MNV virions with addition of ions

The ITC data clearly indicated that metal ions and bile acid influenced NB-5829 binding. A recent study showed that addition of bile acid caused the P dimers to collapse on the shell [32]. To better understand this phenomenon with cations, the cryo-EM structure of virion in complex Mg^2+^ and Ca^2+^ was determined to 4.3 and 4.6 Å resolution, respectively (Fig. S4). Both structures closely resembled each other, where the P dimers were resting on the shell and rotated ∼114° clockwise (Fig. 9), which was similar to bile salt binding [32]. Superposition of the P domain and NB-5829 complex into this cryo-EM structure showed steric clashes between neighbouring Nanobodies (Fig. S6). To investigate this further, we incubated virions with NB-5829 and then added 10 mM MgCl_2._ The cryo-EM structure of this complex was determined to 4.5 Å resolution (Fig. S4). To our surprise, NB-5829 bound to all P dimers and these P dimers were raised off the shell, as observed in the cryo-EM virion and NB-5829 complex structure (Fig. 10). This result indicated that NB-5829 could constrain the effects of added cations and this in turn prevented the P dimers from lowering onto the shell.

**Figure 9.**
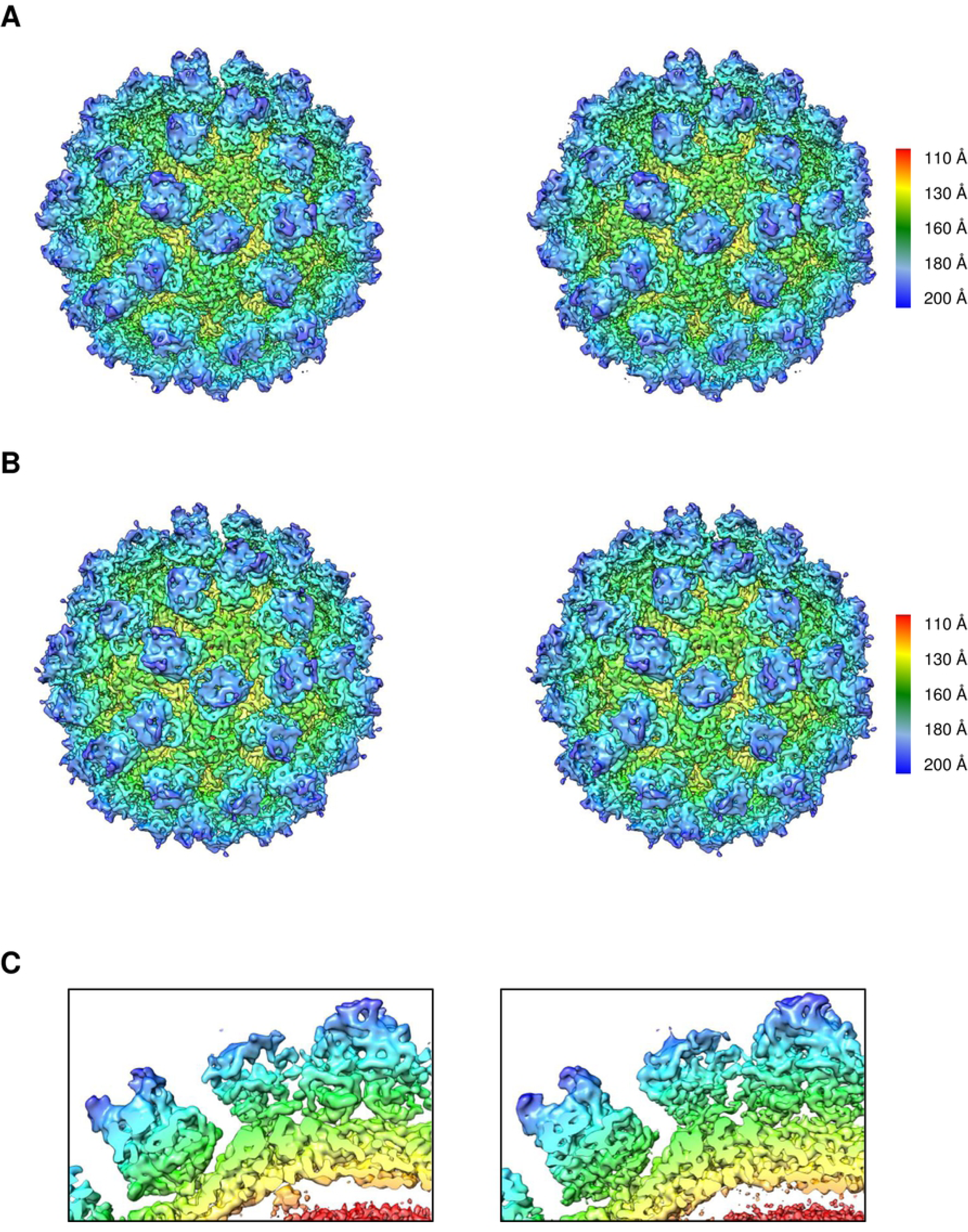
Cryo-EM structures of MNV virion with ions. Icosahedral reconstructions of the virion in with Mg^2+^ and Ca^2+^ were determined to 4.3 and 4.6 Å resolution, respectively. The structures are colored by radius (A) A stereo view of the virion with Mg^2+^. The P domains were rotated ∼114° relative to the apo structure. The P domain was also collapsed on the shell. (B) A stereo view of the virion with Ca**^2+^**. (C) Close-up cutaway view of the S and P domains (left, virion with Mg^2+^; right, virion with Ca**^2+^**) showing the P dimers resting on the shell.

**Figure 10:**
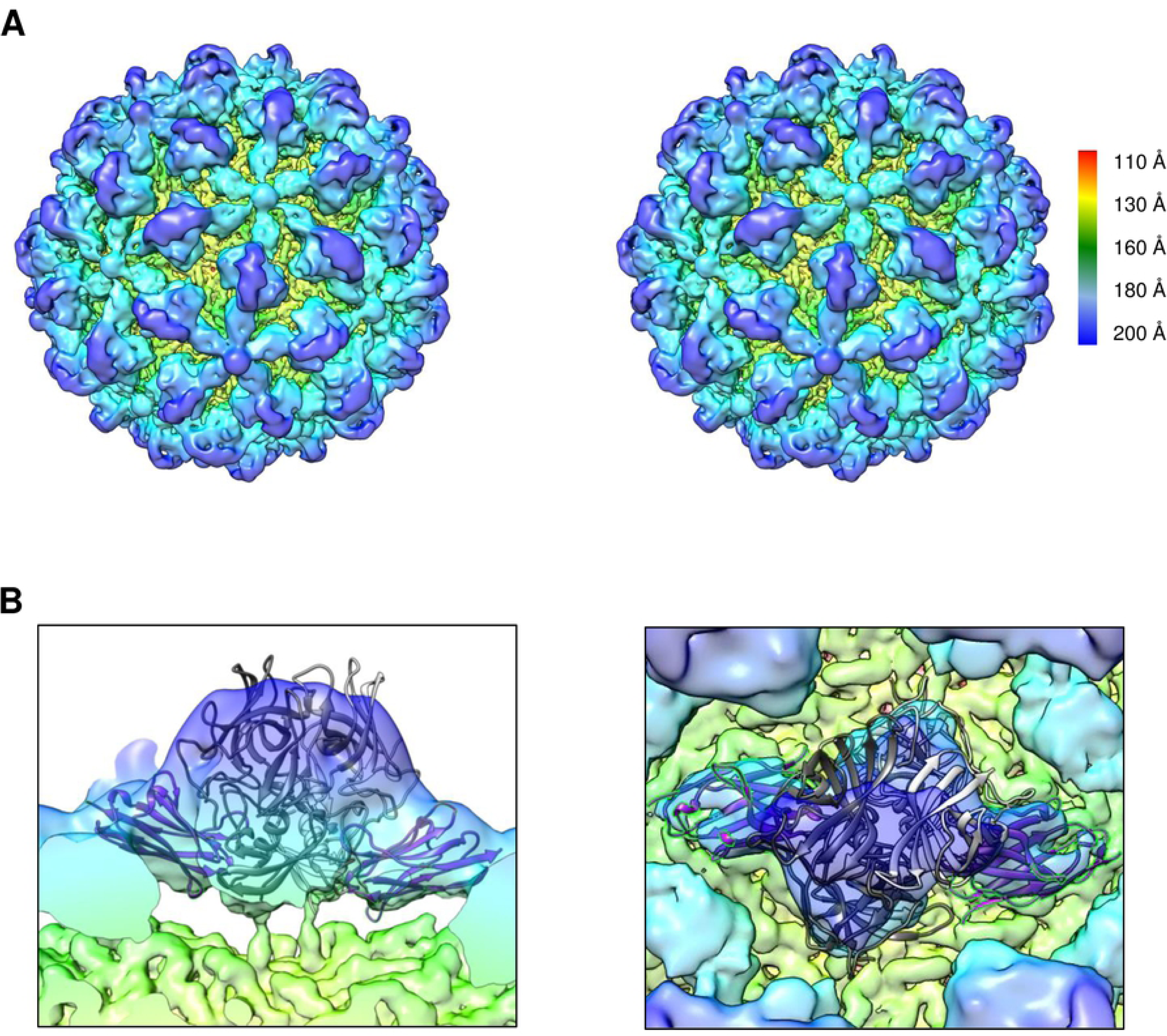
Cryo-EM structure of the MNV virion and NB-5829 complex with Mg^2+^. Icosahedral reconstruction of this mixture was solved to 4.5 Å resolution. The structure is colored by radius (A) A stereo view of the virion complex showed extra density for the Nanobody. Overall, this structure closely resembled the complex structure of MNV and NB-5829 (i.e., without Mg^2+^, see Figure 7). (B) A close-up side view (left panel) and top view (right panel) showing the fitted crystal structure of MNV with NB-5829. The X-ray crystal structure fitted well into the density, indicating that a comparable structural change as with NB-5829 alone occurred.

## DISCUSSION

Norovirus causes a significant number of infections worldwide, with serious health risks to some individuals including the elderly and immunocompromised. The US Centers for Disease Control and Prevention estimates that norovirus is the most common cause of acute gastroenteritis in the United States.

The search for norovirus inhibitors is still in its infancy and there are few reports of antivirals, reviewed in [33]. Most human norovirus capsid antivirals are targeted towards the HBGA pocket [34–38]. In earlier studies, we found several compounds that bind at the HBGA pocket, including human milk oligosaccharides (HMOs) and citrate [39–41]. Other studies have discovered mAbs that partially overlap or inhibit the HBGA pocket [7, 24, 42–44]. Importantly, treatment with mAbs has been linked with a decreased risk of infection and illness [7, 24, 42–44]. We have also analyzed human norovirus-specific Nanobodies that induce structural modifications and blocked VLP attachment to HBGAs [21, 30, 31].

In the current study, we provided proof that MNV-specific Nanobodies were highly capable of neutralizing MNV in cell culture. The top-binding Nanobodies inhibited the receptor-binding site and this in turn blocked particle attachment to cells. Likewise, mAbs A6.2 and 2D3 have an epitope that partially overlaps the receptor site and these mAbs prevented virion attachment to the receptor [27, 28]..Our findings showed that NB-5853 and NB-5867 interacted with P domain residues that bound CD300lf. In fact, NB-5867 and NB-5853 binding induced several structural modifications that were analogous to CD300lf binding.

The dimeric-binding Nanobodies bound to an epitope that connected the S and P domains. This binding event likely blocked a structural modification normally associated with cation and bile acid binding. Recently, divalent cations and bile salts were identified as important co-factors of MNV infection and were able to restore infection [15]. Ca^2+^ and Mg^2+^ ions were essential for the CD300lf binding to the MNV capsid protein, whereas bile acid binding was proposed to slightly increase the receptor binding affinity. Interestingly, bile acid was found to drastically affect the conformation of the MNV particles [32]. In an apo state, the MNV P domain is raised 16-Å off the shell [27]. In the presence of bile acid, the P dimers rotated ∼90° and rested on the shell. Our findings showed that cations induced an equivalent structural modification. Importantly, the dimeric-binding Nanobodies inhibited this structural rearrangement.

We also found that the addition of bile acid or cations dramatically changed the thermodynamic characteristics of Nanobody binding, likely through long-range interactions or solvent rearrangement. Indeed, Ca^2+^ coordination has been shown to impact the stability and structural flexibility of the polyomavirus SV40, allowing virion structural alterations during early steps of infection [45]. Therefore it is tempting to suggest that similar allosteric and long-distance interactions were responsible for the lowering of the MNV P domain onto the shell, perhaps through altered conformations of the hinge region or additional interactions between S and P domains.

Based on these results and previous observations, the following model of MNV entry seems plausible (Fig. 11). In the normal state, MNV virions are in a raised conformation, which can transition to the lowered conformation under the influence of cations or bile. Both raised and lowered conformations might engage the receptor, although the lowered state aids in a higher degree of coordination [32]. Crucially, the lowered conformation is likely required for subsequent, post-receptor attachment steps during cell entry, such as internalization or uncoating. Nanobody binding to a possible conformational switch at the dimeric interface on the side of the P domain might lock the raised conformation and thereby prevented infection. Similar post-binding antibody inhibition mechanisms were described for cytomegalovirus, HIV, respiratory syncytial virus, and chikungunya virus [46–50]. For example, uncoating of rhinovirus could be blocked by antiviral Win compounds that stabilize the capsid by binding to an occluded epitope [51, 52].

**Figure 11.**
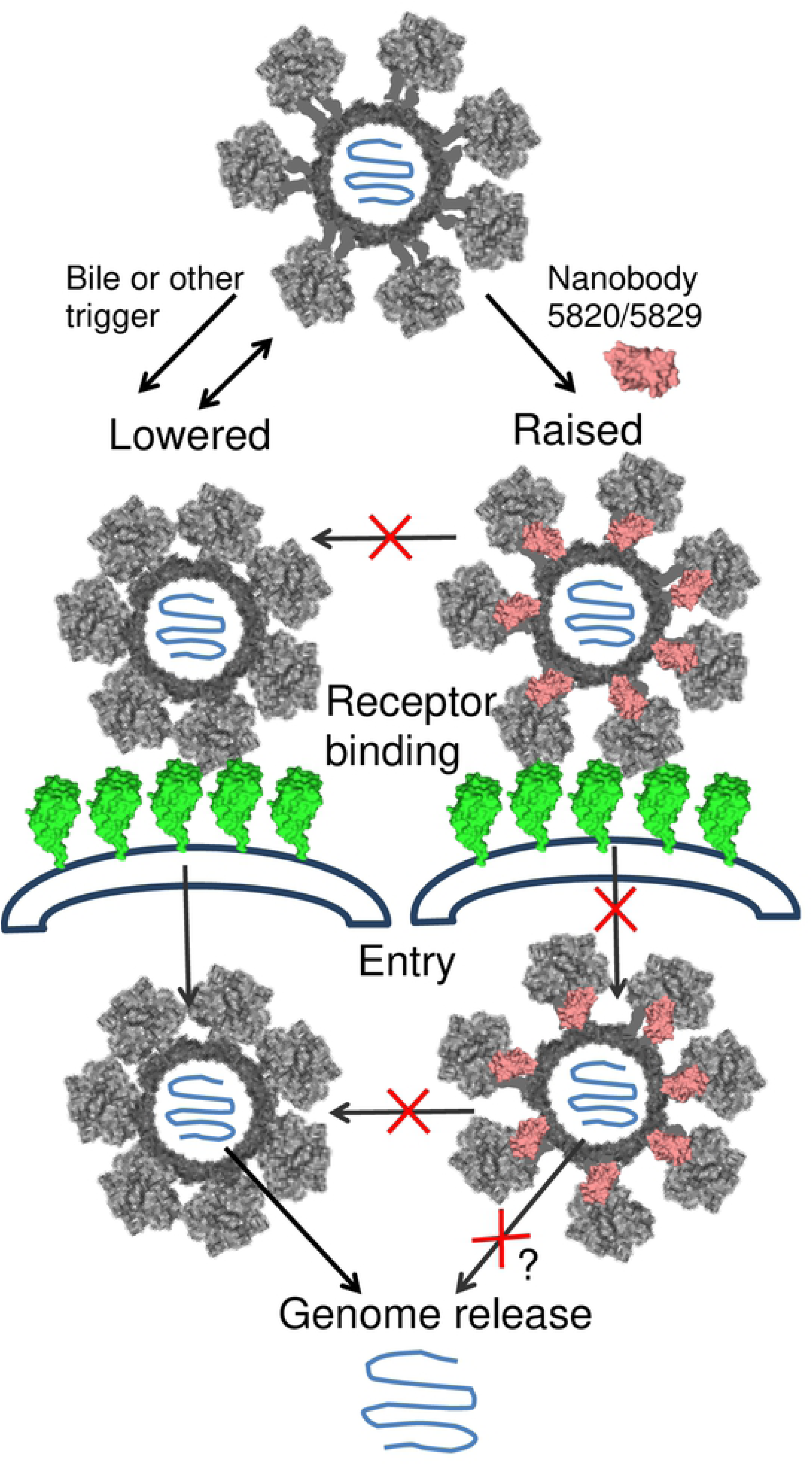
Proposed mechanism of Nanobody 5829 neutralization. In a normal state the capsid is raised, which transitions to a lowered conformation under the influence of external or internal triggers. The raised conformation can be locked by NB-5820 and NB-5829. Both raised and lowered states can engage the receptor, CD300lf. However, the lowered state is required for the subsequent, post-receptor attachment steps during cell entry. Nanobodies binding to a possible conformational switch on the side of the P domain prevents one or several of these steps.

The raised and lowered P dimers have also been observed in human noroviruses and this was thought as a characteristic feature of different genotypes [12, 16, 20]. Therefore, the question remains open whether this feature is uniquely found in MNV or if bile acid or ions induce the lowering of the P dimer for human noroviruses as well. In human norovirus cell culture studies, bile acids were shown to improve infection in some genotypes, which suggested its involvement in the infection process [53]. In other studies, the addition of bile acid could induce VLP binding to HBGAs for a typical non-HBGA-binder genotype [14]. On the other hand, recent studies suggested that the bile acids were not influencing the particles directly, but affected intracellular mechanisms that in turn enhanced infection [54].

Interestingly, our data showed the striking similarity of vulnerable regions on the human norovirus and MNV. In both cases, binding of the Nanobodies to these epitopes alters the normal capsid dynamics and could abort the infection process. In summary, this comprehensive study provided evidence that MNV-specific Nanobodies were highly capable of neutralizing MNV and two different mechanisms of Nanobody-based neutralization were described.

## MATERIALS AND METHODS

### Protein production

MNV-specific Nanobody clones were produced at the VIB Nanobody service facility (Belgium) as previously described [31]. Briefly, a single alpaca was injected with purified MNV virions (MNV.CW1) and a VHH library for MNV-specific Nanobodies was prepared. A total of 125 Nanobody genes were isolated and allocated to 51 distinct families based on CDR sequence diversity. At least one Nanobody per family was selected for cloning and expression. The Nanobody genes were cloned into the pHEN6C vector, expressed in *E. coli* WK6 cells, purified using size exclusion chromatography, and stored in phosphate buffered saline (PBS) or gel filtration buffer (GFB). MNV P domain and the soluble domain of CD300lf (sCD300lf) were produced as described earlier [13].

### Ethics statement

All vaccination experiments were executed by VIB Nanobody service facility following the EU animal welfare legislation and with the approval of the ethics commission of Vrije Universiteit, Brussels, Belgium.

### MNV virion propagation and purification

Murine norovirus (MNV.CW1) was propagated in RAW 264.7 cells as previously described [11]. MNV virions were concentrated using ultracentrifugation and then purified using caesium chloride or sucrose ultracentrifugation gradients [27]. Fractions were checked by negative stain electron microscopy (EM). MNV fractions were pooled, concentrated, and dialyzed into PBS.

### Electron Microscopy

MNV virion morphology was visualized using negative stain EM as previously described [31]. Nanobody and MNV virions were mixed in a 1:1 ratio, incubated for 1 h and applied on EM grids. Grids were stained with 1% uranyl acetate and examined on a Zeiss 900 electron microscope (Zeiss, Oberhofen, Germany) at 50,000× magnification.

### MNV neutralization assays

The MNV titer was determined with a plaque assay as described previously [11, 13]. Virus neutralization by Nanobodies was determined using an attachment plaque assay, an entry plaque assay, and post-entry plaque assay [55]. For the attachment assay, equal plaque forming units of MNV were preincubated with various concentrations of Nanobodies for 1 h at RT and then applied on pre-cooled cell monolayer plates for 3 h at 4°C. Unbound virus was washed twice with ice-cold PBS prior to addition of low melting point (LMP) agarose overlay. For the entry assay, MNV was allowed to attach to the cell monolayer during incubation for 3 h at 4°C. Unbound virus was washed off and serially diluted Nanobodies were then applied for 1 h at 37°C. Plates were washed twice with PBS and covered with a LMP agarose overlay. For the post-entry assay the Nanobodies were added after 1 h at 37°C and were present in the agarose overlay until the end of the assay. The assay was repeated three times to calculate standard deviations.

### ELISA

Nanobody titers were evaluated using a direct ELISA (17). Briefly, microtiter plates were coated with ∼2 µg/ml of MNV virions, washed, and then blocked with skim milk. Nanobodies were first serially diluted, added to the wells, washed, and detected with horseradish peroxidase-conjugated mouse α-Histidine monoclonal antibody. Absorbance was measured at 490 nm (OD_490_) and all experiments were performed in triplicate.

### Isothermal Titration Calorimetry (ITC) measurements

ITC experiments were performed using an ITC-200 (Malvern, UK). Samples were dialyzed into the identical PBS buffer and filtered prior titration experiments. Titrations were performed at 25°C by injecting consecutive (1-2 µl) aliquots of Nanobodies (100-150 µM) into MNV P domain (10-20 µM) in 120 second intervals. The binding data was corrected for the heat of dilution and fitted to a one-site binding model. For the ITC measurements with bile acid and calcium, the P domain was pre-mixed with 50 µM GCDCA or 5 mM CaCl_2_. Titrations with Nanobodies were then performed as above. For competitive ITC experiments, 200 µM sCD300lf was titrated into MNV P domain premixed with GCDCA and CaCl_2_. The sample cell contents were then used for the subsequent titration with Nanobody. For the measurements with bile, 100 µM GCDCA was mixed with the MNV virion, and then Nanobodies were titrated as above. All experiments were performed twice.

### P domain and Nanobody complex purification and crystallization

For each complex, the MNV (CW3) P domain and Nanobody were mixed in a 1:1.5 molar ratio and incubated overnight at 4°C. The complex was purified by size exclusion chromatography using a Superdex-200 column and the peak shift observed in the UV elution profile clearly indicated the presence of the P domain/Nanobody complex in solution. The mixture was then concentrated to ∼4 to 6 mg/ml. When required for the structural study, 100 µM of bile acid (GCDCA) were added at this stage to the concentrated mixture. Crystallization conditions were screened using the sitting-drop vapor diffusion method and optimized using the hanging drop method. P domain NB-5820: 25% PEG3000, and 0.1 M Tris (pH 8.5); P domain NB-5829: 0.2 M sodium chloride, 0.1 M phosphate-citrate (pH 4.2), and 20% PEG8000; P domain NB-5853: 20% PEG 3000 and 0.1 M sodium acetate; and P domain NB5867: 20% PEG 6000 and 0.1 M Citric acid (pH 5). Crystals were soaked in a cryoprotectant solution containing the same mother liquor with an addition of 30% 1,2-ethandiol, then flash frozen at 100K.

### Data collection, structure solution, and refinement

X-ray diffraction data were collected at the European Synchrotron Radiation Facility, France at beamlines ID29, ID30A-3, and ID30B, and processed with XDS [56]. The structures were solved by molecular replacement in PHASER *Phaser-MR* [57] using the monomeric structure of the previously solved murine norovirus P domain (PDB code 3LQ6). Alternate cycles of manual model rebuilding with COOT [58] and refinement with the REFMAC5 (CCP4 program suite) [59, 60] were used. Final structures have been validated with Molprobity [61] and wwPDB Validation Server at (https://validate.wwpdb.org). Co-ordinates of the final structures have been deposited in the Protein Data Bank (P domain in complex with NB-5853: 6XW6; P domain in complex with NB-5867: 6XW4; P domain in complex with NB-5820: 6XW5; and P domain in complex with NB-5829: 6XW7).

### Cryo-EM of MNV virions

Purified MNV virions (1 mg/ml) were diluted in PBS. These were loaded onto freshly glow discharged Quantifoil 0.6/1 grids and vitrified using a Mark IV vitrobot, held at 12°C and 100% humidity. After blotting the samples for 20 s, the samples were plunge frozen in liquid ethane and stored in liquid nitrogen. Data collection was performed on a Titan Krios, operated at 300 keV, equipped with a K3 direct electron detector. Movies were collected at 64,000× magnification, corresponding to a pixel size of 1,368 Å/px. The movies, containing 15 frames, were corrected for drift using MotionCor2 [62] and defocus estimation was done with Ctffind4.1 [57], as implemented in Relion 3 [63]. All further processing steps were performed in Relion 3. Apo particles were picked automatically from 1,588 micrographs. Following 2D and 3D classification, a subset of 21,424 particles was used for icosahedral reconstruction, leading to a resolution of 4.6 Å, as determined using Fourier shell correlation (FSC) with a cut-off at 0.143. The structure was deposited in the EMDB (EMD-10596).

### Cryo-EM of virion and NB-5829 complex

MNV virions (1 mg/ml) were incubated with NB-5829 (1 mg/ml) for 30 min at RT before vitrification. Particles were prepared and processed as described above. For this complex, 32,612 particles were picked from 3,599 micrographs. Refinement of the particles resulted in a structure at 4.7 Å resolution (FSC cut-off at 0.143). The density was deposited in the EMDB under accession code EMD-10597. Focused reconstruction of this complex was also performed as described earlier [64]. Briefly, the symmetry of the dataset was expanded following icosahedral refinement, so that each particle was assigned 60 orientations that were matching the repeated views of the icosahedral particle. With this expanded dataset, 3D classification without orientation refinement was done to resolve the asymmetric differences within the single VP1 dimers. To include only the VP1 dimers in the classification, a cylindrical mask was prepared in SPIDER [65] and positioned in UCSF Chimera [58], to only include the single capsomers.

### Cryo-EM of virion and cation complexes

MNV virions (1 mg/ml) were incubated with 10 mM MgCl_2_ or CaCl_2_ prior to vitrification. Particles were prepared and processed as described above. For MNV and Mg^2+^, 2,783 particles from 611 micrographs were used for final reconstruction and resulted in a resolution of 4.3 Å (FSC cut-off at 0.143). For MNV and Ca^2+^, 2,637 particles were picked from 303 micrographs that lead to a final resolution of 4.6 Å (FSC cut-off at 0.143). For the complex of MNV/NB-5829/Mg^2+^, the virions were first incubated with NB-5829 for 30 min at RT, and then 10 mM MgCl_2_ for 3 h at RT. A total of 7,386 particles from 293 micrographs revealed a structure of 4.5 Å (FSC cut-off at 0.143). The structures have been deposited in the EMDB as EMD-10598 (MNV with Mg^2+^), EMD-10599 (MNV in complex with Ca^2+^), and EMD-10600 (MNV with NB-5829 and Mg^2+^).

## ACKNOWLEDGEMENTS

We thank David Bhella and Götz Hofhaus for assistance with Cryo-EM data collection and structural refinements; and Benedikt Wimmer for setting up the cryo-EM software. We acknowledge the excellence cluster CellNetworks (Cryo-EM network) of the University of Heidelberg for cryo-EM data collection, the EM core facility at DKFZ, Baden-Württemberg High Performance Cluster (bwHPC), and the data storage service SDS@hd. We are grateful to the Protein Crystallization Platform facility of BZH-Heidelberg University. We also thank the European Synchrotron Radiation Facility and staff (France) for use and help at beamlines ID29, ID30A-3, and ID30B.

## SUPPLEMENTARY FIGURE LEGEND

**Figure S1: Neutralization using Nanobodies in cell culture.** (A) The neutralization capacity of Nanobodies was analysed in an attachment assay. Nanobodies were tested at 20 μg/ml. In total, 38 of 58 Nanobodies showed inhibition greater than 70%. All experiments were performed in triplicates and standard deviation is shown. (B) MNV neutralization was tested with serially diluted (1:4 or 1:10) Nanobodies with a starting concentration of 2 μg/ml. Nanobodies had IC_50_ values ranged between 0.03 to 1.6 μg/ml. All experiments were performed in triplicates and standard deviation is shown.

**Figure S2: Binding footprints of Nanobodies to P domain.** As comparison, the binding footprints of the receptor CD300lf (green) is shown. NB-5853 (blue) and NB-5867 (purple) exhibit closely overlapping binding footprints with CD300lf. NB-5820 (salmon) and NB-5829 (pink) have epitopes located on the side of the P domain and interact with both P domain monomers.

**Figure S3: Negative-stain EM of treated.** Untreated virions were well dispersed and showed little aggregation. After incubation with NB-5820 and NB5829, the particles appeared as the untreated particles. Incubation with NB-5853 and NB5867 led to partial aggregation.

**Figure S4: Cryo-EM analysis of virion.** Shown are raw micrographs (left column) for apo virion, virion/NB-5829, virion/MgCl_2_, virion/CaCl_2_, and virion/NB-5829/MgCl_2_. The center column shows the central section of the structure. The FSC curves (right column) indicated resolutions of 4.6 Å for the virion, 4.7 Å for virion/NB-5829, 4.3 Å for virion/MgCl_2_, 4.6 Å for virion/CaCl_2_, and 4.5 Å for virion/NB-5829/MgCl_2_.

**Figure S5: Cryo-EM structure of the apo virion.** (A) A stereo view of the apo virion colored by radius. The P dimers (cyan/blue) protruded from the shell (yellow/green). (B) The X-ray crystal structure of the P domain NB-5829 complex (P domain: blue and NB-5829: purple) was fitted into the apo virion structure. In this model, the Nanobodies clashed with adjacent Nanobodies at the 3-fold and 5-fold icosahedral axes.

**Figure S6: Structural restraints of the lowered P dimers.** The X-ray crystal structure of the P domain complex was fitted into the cryo-EM structure of virion/NB-5829/MgCl_2_. In this model, the Nanobodies clashed with adjacent Nanobodies at both 3-fold and 5-fold icosahedral axes.

## SUPPLEMENTARY TABLE LEGEND

**Table S1: Summary of characterized Nanobodies.** Nanobodies were sorted by family- and Nanobody number. The table displays the cut-off value for binding as determined by ELISA; the IC_50_ value as determined by attachment assay; the binding site of the Nanobody; and Nanobody aggregation ability as observed using EM. The table shows the K_d_, dH, and -TdS values for eight Nanobodies tested using ITC.

**Table S2: Summary of X-ray crystallography structures.** The table displays data collection and refinement details.

**Table S3: Binding interactions between P domain and Nanobodies.** Both top binding Nanobodies only interact with one monomer per P dimer. The dimeric-binding Nanobodies interact with both subunits of the P domain dimer. Mentioned are hydrophobic, H bond and water bridge mediated interactions.

